# Stability of ligand-induced protein conformation influences affinity in maltose-binding protein

**DOI:** 10.1101/2021.02.26.433038

**Authors:** Marco van den Noort, Marijn de Boer, Bert Poolman

## Abstract

Our understanding of what determines ligand affinity of proteins is poor, even with high-resolution structures available. Both the non-covalent ligand-protein interactions and the relative free energies of available conformations contribute to the affinity of a protein for a ligand. Distant, non-binding site residues can influence the ligand affinity by altering the free energy difference between a ligand-free and ligand-bound conformation. Our hypothesis is that when different ligands induce distinct ligand-bound conformations, it should be possible to tweak their affinities by changing the free energies of the available conformations. We tested this idea for the maltose-binding protein (MPB) from *Escherichia coli*. We used singlemolecule Förster resonance energy transfer (smFRET) to distinguish several unique ligand-bound conformations of MBP. We engineered mutations, distant from the binding site, to affect the stabilities of different ligand-bound conformations. We show that ligand affinity can indeed be altered in a conformation-dependent manner. Our studies provide a framework for the tuning of ligand affinity, apart from modifying binding site residues.

## INTRODUCTION

Insight in the principles behind ligand affinity of proteins is fundamental to all biological processes. From an evolutionary perspective, selection pressure can lead to adjustments in ligand affinity of transporters, receptors and enzymes, depending on the need and availability of a given ligand or substrate (1–3). Approaches to tweak the ligand affinity of a protein are popular for applications in biocatalysis. Retrospective methodologies, such as ancestral protein reconstruction and directed evolution, have been introduced to speed up the engineering of proteins (1, 3, 4). Information from these studies and thermodynamic models are needed to rationalize the findings and make the redesign of proteins with new or improved functions more predictive. It is tempting to reduce the concept of ligand affinity to only non-covalent protein-ligand interactions. However, one then narrows the possibilities of protein engineering, which is also evident from screens based on directed evolution (3–6).

Advances in the fields of electron paramagnetic resonance (EPR) (7–9), molecular dynamics (MD) (5, 8–13) and single-molecule spectroscopy (14–19) allow visualization of protein conformational dynamics on a femtosecond to even second time scales. This led to increased appreciation of the concept that different protein conformations are a determinant of ligand affinity and catalytic activity. Examples are the acyltransferase enzyme LovD (5), adenylate kinase (18), epoxide hydrolase (13), the substrate-binding domains (SBDs) of the ATP-binding cassette (ABC) transporter GlnPQ (19) and substrate-binding protein (SBP) derived cyclohexadienyl dehydratases (9).

Also for the maltose-binding protein (MBP) from *Escherichia coli* it has been shown that conformational dynamics is a determinant of ligand affinity (20–26). MBP belongs to a very large class of SBPs associated with an importer type of ABC transporter or ligand receptor or ion channel (15, 27, 28). MBP, also known as MalE, is the best-studied SBP and consists of two lobes, which are connected by a flexible hinge region. MBP samples a single (open) conformation in the absence of ligands, which does not undergo significant conformational changes on the millisecond to second timescale (20, 29). Ligand binding occurs via an ‘induced fit’ mechanism, wherein the ligand binds to the open conformation and induces structural rearrangements in both lobes and the hinge to form a closed state (20, 29). The closed conformation is structurally significantly different from the open conformation. Mutations and regio-specific synthetic antibodies that alter the free energy difference between the two states have been shown to tune the affinity for maltose, while the binding pocket residues are unaltered (20–22, 25, 26).

The simple model of ligand binding is based on the binding of maltose and one closed conformation. However, MBP is very promiscuous and able to bind a wide range of maltose-derivatives with affinities in the sub-μmolar and μmolar range (30, 31). Additionally, and contrary to what X-ray structures of MBP suggest, the protein doesn’t merely use the open and one closed conformation to bind the different ligands (32–35). Instead, MBP samples a range of different ligand-specific closed conformations (29, 36, 37). Part of the promiscuity of MBP can be attributed to the architecture of its binding pocket (38). However, we now show that affinity can be tuned by addressing the stability of the distinct closed conformations of the protein.

In this work, we consider that the dissociation constant (K_D_) of ligand binding by the protein is given by:

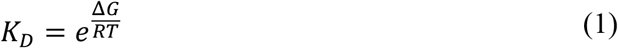

where Δ*G* is the overall free energy of ligand binding, *R* is the gas constant and *T* is the absolute temperature. Thermodynamics dictates that ΔG can be expressed as a sum of the conformational free energies, Δ*G_conf_* = *G_C_* — *G_O_*, where G_C_ and *G_O_* are the free energies of the open and closed conformations of the apo protein, respectively; and the intrinsic affinity of the binding site for the ligand, Δ*G*_bind_ = *G_CL_ — μ_0_ — G_C_*, where G_CL_ is the free energy of the closed-liganded conformation and μ_0_ is the standard chemical potential (**Fig. 1**) (details in Material and Methods):

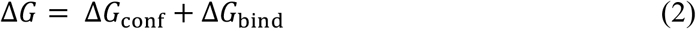

**FIGURE 1.**
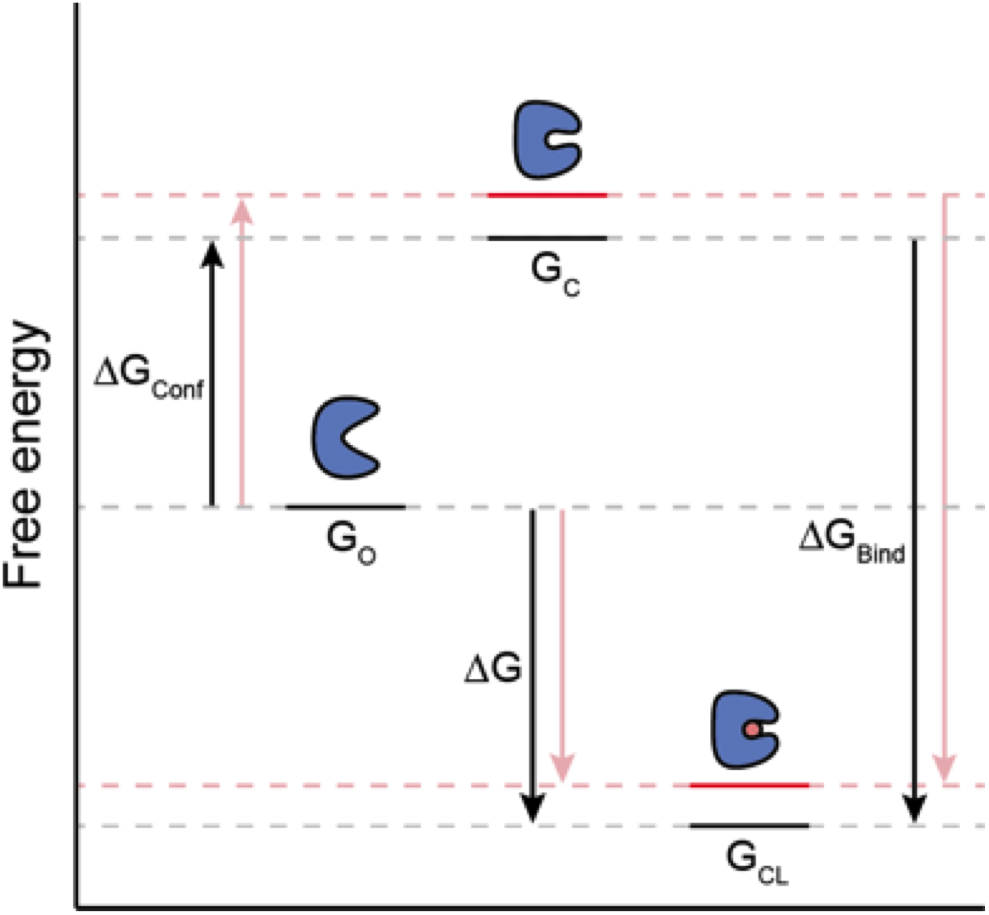
A model of the relationship between ΔG_conf_ and Δ*G*_bind_ with ΔG before (black) and after a mutation that alters the free energy of the ligand-bound conformation (red). The model considers an open state in the absence of ligands (G_O_), a theoretical ligand-free closed state (G_C_) and a ligand-bound closed state (G_CL_). The amount of change in G_C_ and G_CL_ after the mutation is equal if the considered closed conformation is the same. Hence, this type of mutation only affects ΔG_conf_ and the decrease in ΔG is equal to the increase in ΔG_conf_.

An important insight from **Eq. 2** is that different ligands that each induce the same closed conformation share the same ΔG_conf_ but not necessarily the same Δ*G*_bind_, whereas ligands that induce different closed conformations do not necessarily share either the same ΔG_conf_ or Δ*G*_bind_. Following from **Eq. 1** and **2**, when the ΔG_conf_ is changed by altering G_C_ (and G_CL_ accordingly), then the Δ*G* is altered equally for every ligand that induces the same closed conformation and is altered differently when the ligands do not induce the same closed conformation (**Fig. 1**). Following from **Eq. 1**, an equal change in ΔG will be manifested by a change in the K_D_ by an equal factor (**Eq. 12**).

To experimentally confirm this hypothesis, we identified and mutated residues that likely have different molecular environments in the different closed conformations. These were analysed by means of single-molecule Föster resonance energy transfer (smFRET) and bulk intrinsic protein fluorescence measurements to assess the different conformations and determine the K_D_ values, respectively.

## RESULTS

### Correlation between ligand affinity and conformational stability in MBP

In general, a ligand that binds to an SBP induces a relative twist and closure of the two lobes (27, 29). In the case of MBP, the degree of closure is ligand dependent (29, 36, 37). To assess the different degrees of closure, a thiol-reactive, maleimide-based donor and acceptor fluorophore were linked to engineered cysteines at position 36 and 352 in the protein sequence (29). These positions are located on top of the two MBP lobes and exhibit a relative distance change of several angstroms upon binding of a ligand (**Fig. 2A**). We examined the conformations of freely diffusing and labeled proteins, using a confocal microscope with alternating laser excitation (ALEX)(39). The labeled proteins were monitored upon transition through the laser spot, where donor- and acceptor-based fluorescent signals are triggered after donor excitation (**Fig. 2B**). Their relative fluorescent intensities allow for an apparent FRET efficiency (E*) estimation of each recorded MBP molecule, which relates back to the inter-fluorophore distance. The subsequent excitation of the acceptor is used to determine the stoichiometry (S) of the fluorophores. MBP molecules with an S of ~0.5 exhibit an E* that is dependent on the relative closure of the two lobes (**Fig. 2B**). The other molecules with a stoichiometry of near 0 or 1 were filtered out, since these molecules possess only acceptor or donor fluorophores, respectively.

**FIGURE 2.**
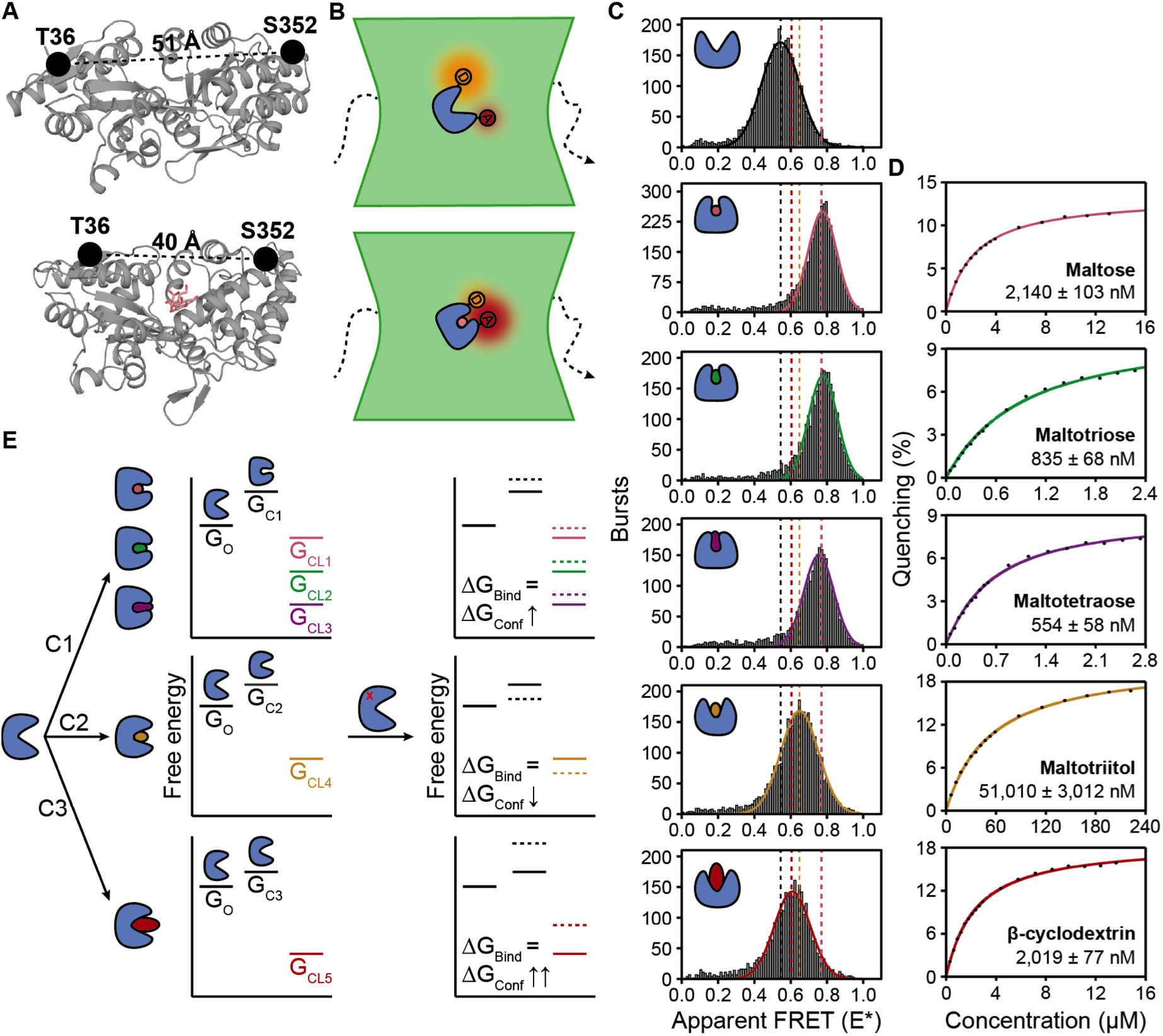
Closed conformation-dependent ligand affinity tuning. (A) Crystal structures of MBP in the absence of ligands (PDB: 1OMP) and in the presence of maltose (PDB: 1ANF). Black dots and lines indicate the fluorophore labeling positions and the distance change upon maltose binding, respectively. (B) A schematic representation of a labeled MBP molecule that diffuses through the confocal volume of a laser beam during a solution-based smFRET measurement. (C) Histograms of selected bursts in ALEX measurements with labeled MBP in the absence of ligand or from top to bottom in the presence of either 1 mM maltose (pink), maltotriose (green), maltotetraose (purple), maltotriitol (yellow) or β-cyclodextrin (red). Solid lines indicate best fit of a single Gaussian distribution. The means of the Gaussian fits are indicated by dotted lines. The dotted lines in pink represent a general mean for the conditions with maltose, maltotriose and maltotetraose. The numbers of included bursts per plot are given in **Table S3**. (D) Examples of intrinsic protein fluorescence-based ligand affinity measurements, using wildtype MBP. Colored lines represent the best fit of the data, using **Eq. 16** in the Materials & Methods section. Numbers in the graph indicate the mean K_D_ ± two times the error of the mean (**Table S2**). (E) Exemplary model to illustrate the theory of closed conformation-dependent ligand affinity tuning. Solid lines indicate theoretical free energy states in wildtype and dotted lines indicate changed states after introducing non-binding pocket mutations (red cross). The model considers one open state in the absence of ligands (G_O_), three theoretical ligand-free closed states (G_C1-3_) and 5 ligand-bound closed states (G_CL1-5_). The amount of change in G_c_ and G_CL_ after a mutation is equal if the considered closed conformation is the same, because this type of mutation is considered to only affect Δ*G_conf_*. Ligand-associated colors in (D) and (E) are the same as in (C).

MBP in the absence of ligand has a mean E* of ~0.6, representing a Gaussian distribution around one conformation (**Fig. 2C**). In the presence of maltose, maltotriose, maltotetraose, maltotriitol or β-cyclodextrin, three different closed conformations could be distinguished: 1) the fully closed, maltose/maltotriose/maltotetraose bound conformation (mean E* ~0.81); 2) the partially closed, maltotriitol bound conformation (mean E* ~0.69); and 3) the partially closed, β-cyclodextrin bound conformation (mean E* ~0.65)(**Fig. 2C**). This is in good agreement with earlier observations (29).

Next, we assessed the affinity of (unlabeled) MBP for the different ligands by means of intrinsic protein fluorescence. MBP excited with light of 280 nm yields fluorescence with an emission maximum around 348 nm, mainly due to the presence of aromatic amino acids. Upon ligand binding, the internal fluorescence is quenched by 5-10%, in concert with a ligand-dependent shift of the emission spectrum (40). This provides an elegant opportunity to accurately determine the dissociation constant (K_D_) for the different ligands, by measuring the change in fluorescence during titration of a ligand and using a protein concentration of only 0.2 μM. Hence, we were able to determine the dissociation constants with an error of just 1-6% of the K_D_ (n ≥ 3) (**Fig. 2D**)(**Table S2**). All K_D_ values were in the (sub-)μmolar range, which is in good agreement with previously reported affinities (21, 24, 25, 30, 32, 41).

Together, the availability of different ligand-induced conformations and the ability to accurately measure affinity allows us to test the hypothesis that the free energies of the ligand-induced closed states can set the ligand affinities. To do so, we defined three different (theoretical) ligand-free closed conformations each with a unique free energy state (**Fig. 2E**):

1. G_C1_ is the free energy of the closed conformation for the binding of maltose/maltotriose/maltotetraose
2. G_C2_ is the free energy of the closed conformation for the binding of maltotriitol
3. G_C3_ is the free energy of the closed conformation for the binding β-cyclodextrin

Note that, in the case of MBP, these conformations are not sampled in solution without the presence of a ligand and are therefore regarded as high energy states compared to the free energy of the open conformation (GO) (20, 29). In the case of closed-liganded conformations we defined five unique free energy states (G_CL1-5_), because these free energies both depend on the free energy of the closed conformation and the protein-ligand interactions (**Fig. 2E**). Now, if a non-binding pocket mutation would differentially change the free energy of the distinct apo-closed conformations while leaving the ligand-protein interactions unaffected, this would be manifested as an equal change in Δ*G_conf_* (ΔΔ*G_conf_*; see **Eq. 13**) for the binding of maltose/maltotriose/maltotetraose, and two different ΔΔ*G_conf_* values for the binding of maltotriitol and β-cyclodextrin, while the Δ*G*_bind_ remains unaltered in all cases. Given **Eq. 2**, such mutations would induce a similar change in ΔG (ΔΔG) as in wildtype MBP for maltose, maltotriose and maltotetraose binding, whereas the ΔΔG for both maltotriitol and β-cyclodextrin binding would change differently. A theoretical example is depicted in **Fig. 2E**. An equal ΔΔG would be manifested by a change in K_D_ with an equal factor (**Eq. 12**).

### Affinity changes by mutations distant from the binding pocket: Closed conformation-dependent changes in ΔG

To test the theoretical framework, we mutated amino acids at sites in MBP that were distant from the binding pocket. Candidate positions were chosen, based on two criteria: (i) the amino acid is not conserved in related SBPs; and (ii) the amino acid makes unique interactions in the maltose-bound crystal structure compared to the ligand free structure (analyzed by the RING 2.0 web server (42)). We ended up with five different positions in MBP: Ser-233, Pro-298, Ile-317, Asn-332 and Pro-334 (**Fig. 3A,B**). The amino acids Ser-233, Pro-298 and Ile-317 are found in the same region. Here, the residues at both lobes move towards each other upon maltose binding. Asn-332 and Pro-334 are located at the opposite site of MBP, but also here the residues at the opposite lobes display movements towards each other upon maltose binding (**Fig. 3A,B**). It has been proposed that Asn-332 is important in the stabilization of the maltose-bound closed conformation, that is, by making extra H-bonds with the backbones of Ala-96 and Gly-68 (24). Thereby, it bridges the two different lobes of MalE and stabilizes the maltose-bound closed conformation.

**FIGURE 3.**
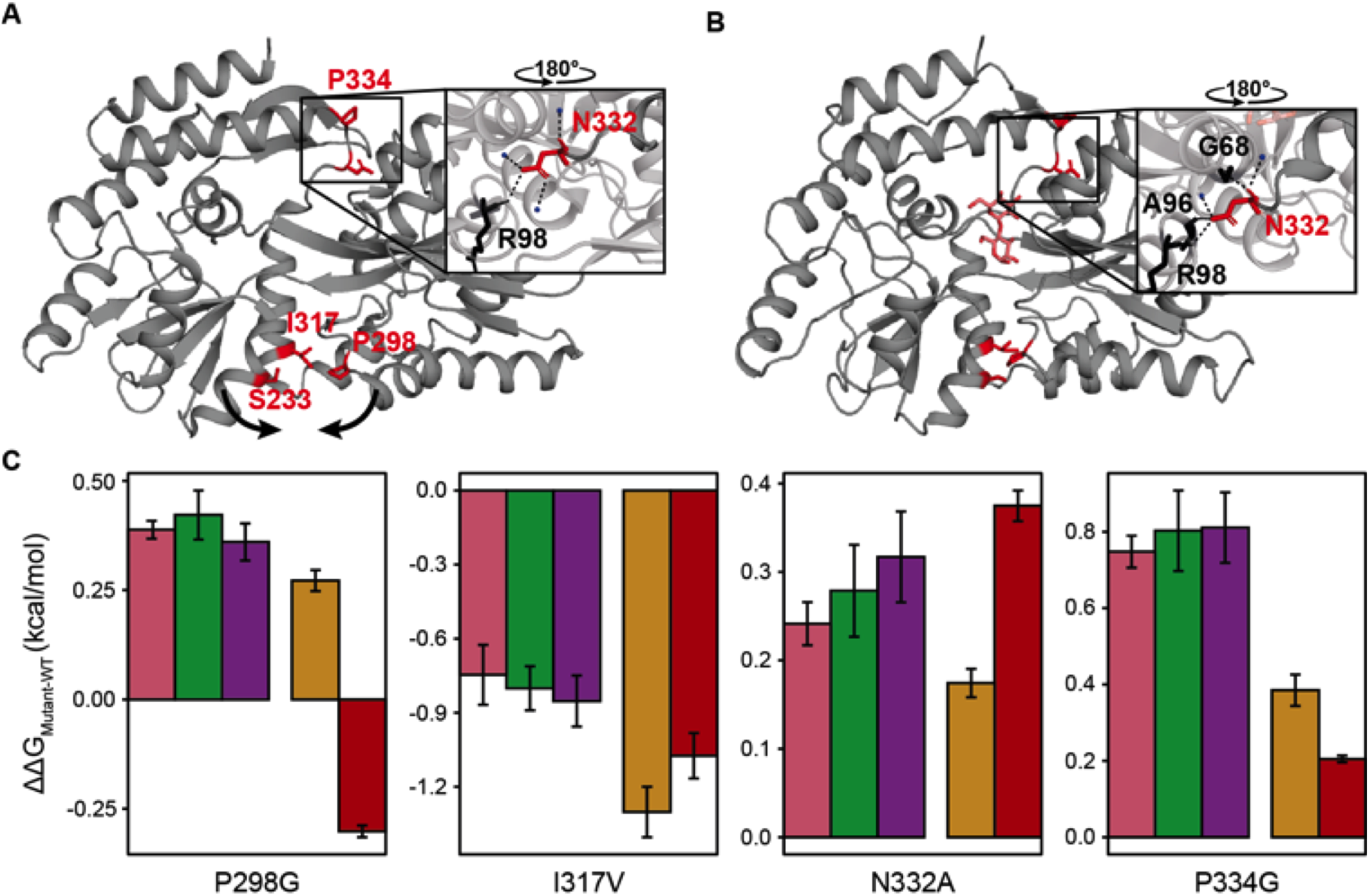
Mutagenesis of sites in MBP distant from the binding pocket reveals closed conformation-dependent changes in ΔG. (A,B) Crystal structures of MBP in the absence of ligand (PDB: 1OMP) and in the presence of maltose (PDB: 1ANF). Mutated residues are indicated in red, and residues that interact with N332 in black. (C) ΔΔG_WT-mutant_ between the open and ligand-bound MBP conformation for maltose (pink), maltotriose (green), maltotetraose (purple), maltotriitol (yellow) and β-cyclodextrin (red) as ligand. Error bars indicate two times the standard error of the mean.

Next, we constructed the following mutants: P298G, I317V, N332A and P334G. With smFRET we confirmed that maltose, maltotriose and maltotetraose induces a similar shift in E*, while maltotriitol and β-cyclodextrin both induced a unique and smaller shift in E* (**Fig. S1**). The exception is the maltotetraose induced E* of P334G, which was different from the values in the presence of maltose or maltotriose. In the case of P298G in the presence of maltotriitol or β-cyclodextrin, there is a difference in E* of only ~0.015, implying that the two conformations are more or less similar or that the labels and the labeling positions do not resolve the two conformations. Nonetheless, P298G is still expected to show a similar ΔΔG for maltose, maltotriose and maltotetraose.

In line with our theory, all four mutants had a similar ΔΔG for the binding of maltose, maltotriose and maltotetraose, and a different ΔΔG for the binding of maltotriitol and β-cyclodextrin (**Fig. 3C**). In the case of P298G it was even possible to tune the affinity to an extent that β-cyclodextrin was bound with a higher affinity (reflected by a negative ΔΔG), whereas the other four ligands were bound less strongly. Unfortunately, in the case of N332A the ΔΔG of the different ligands was too small to unambiguously conclude that the ΔΔGs for maltotetraose and β-cyclodextrin binding are different.

### Affinity changes by mutations distant from the binding pocket: New ligand-induced closed conformations

At position Ser-233 we introduced a harsh substitution to tryptophan, because a mutation to glycine did not exhibit a change in affinity. Unexpectedly, the ΔΔG_WT-S233W_ between the open and ligand-bound MBP conformation for maltose, maltotriose and maltotetraose was not equal (**Fig. 4A)**. S233W displays different closed conformations for the three ligands, *i.e*. based on our smFRET measurements (**Fig. 4B**). In support of the conclusions from the smFRET measurements, binding of maltotetraose elicited a maximal quenching of only 6-7% at 330 nm, which compares to 9-10% for the wildtype protein, and this is indicative for different binding mode for maltotetraose. Consequently, we cannot assume an equal G_c_ or ΔG_conf_ for the three situations anymore. This implies that in S233W the ΔΔG between the open and ligand-bound MBP conformation is not equal for the three ligands.

**FIGURE 4.**
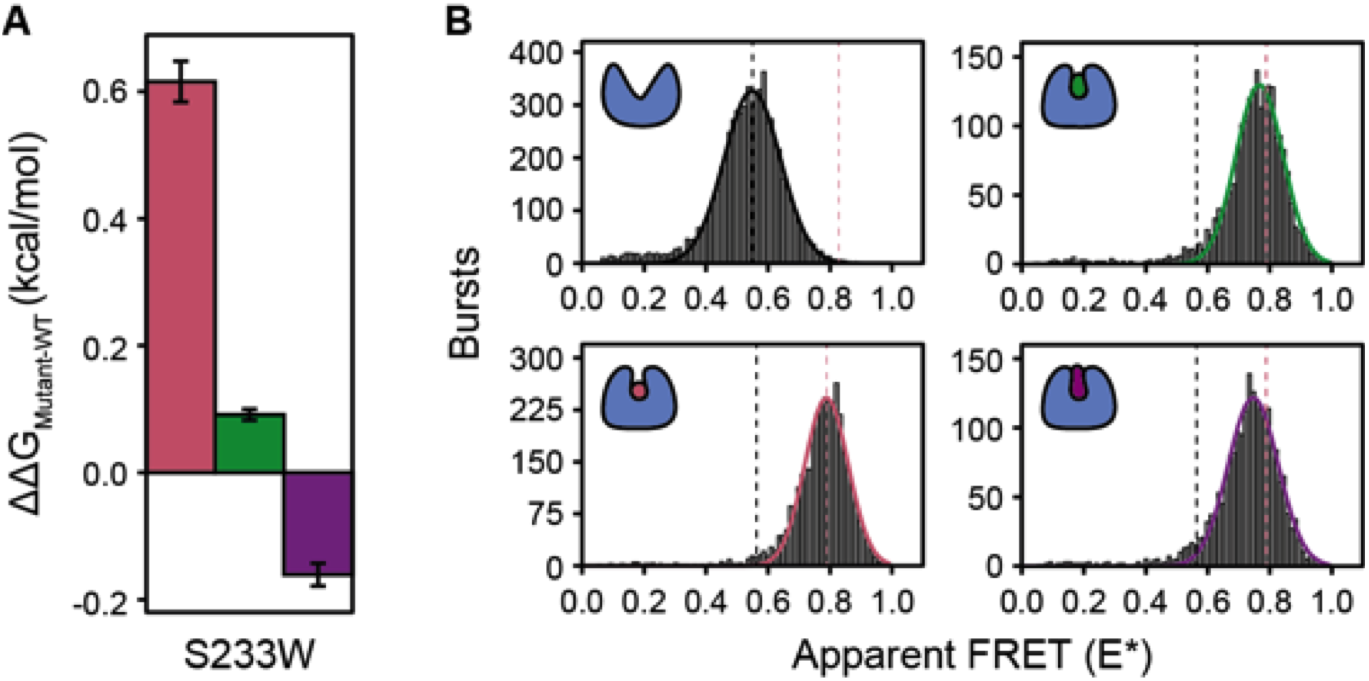
Maltose (pink), maltotriose (green) and maltotetraose (purple) induce different closed conformations in MBP-S233W. (A) ΔΔG_WT-mutant_ between the open and ligand-bound MBP conformation. Error bars indicate two times the standard error of the mean. (B) Histograms of selected bursts after ALEX measurements with MBP-S233W in the absence of ligand and in the presence of either 1 mM maltose, maltotriose or maltotetraose. Solid lines indicate the best fit of a single Gaussian distribution. The means of the ligand-free and maltose distributions are represented as dotted lines. The numbers of included bursts per plot are given in **Table S3**.

### Affinity changes by mutations at the periphery of the binding pocket: Closed conformation-independent changes in ΔG

In line with our hypothesis, mutations that affect the direct interactions with the ligand in and around the binding pocket are expected to primarily alter Δ*G*_bind_ instead of ΔG_conf_. Consequently, in such mutants the ΔΔG between the open and ligand-bound MBP conformations would no longer correlate with the ligand-induced closed conformation anymore (for an example, see **Fig. 5A**). Accordingly, we should be able to tune the affinity of ligands that induce a similar ligand-bound conformation, like maltose, maltotriose and maltotetraose, by mutating residues that interact differently with these three ligands. To test this, we changed Glu-44 and Glu-153 to alanine. Both residues reside at the periphery of the binding pocket and bond differently to maltose, maltotriose and maltotetraose (**Fig. 5B**).

**FIGURE 5.**
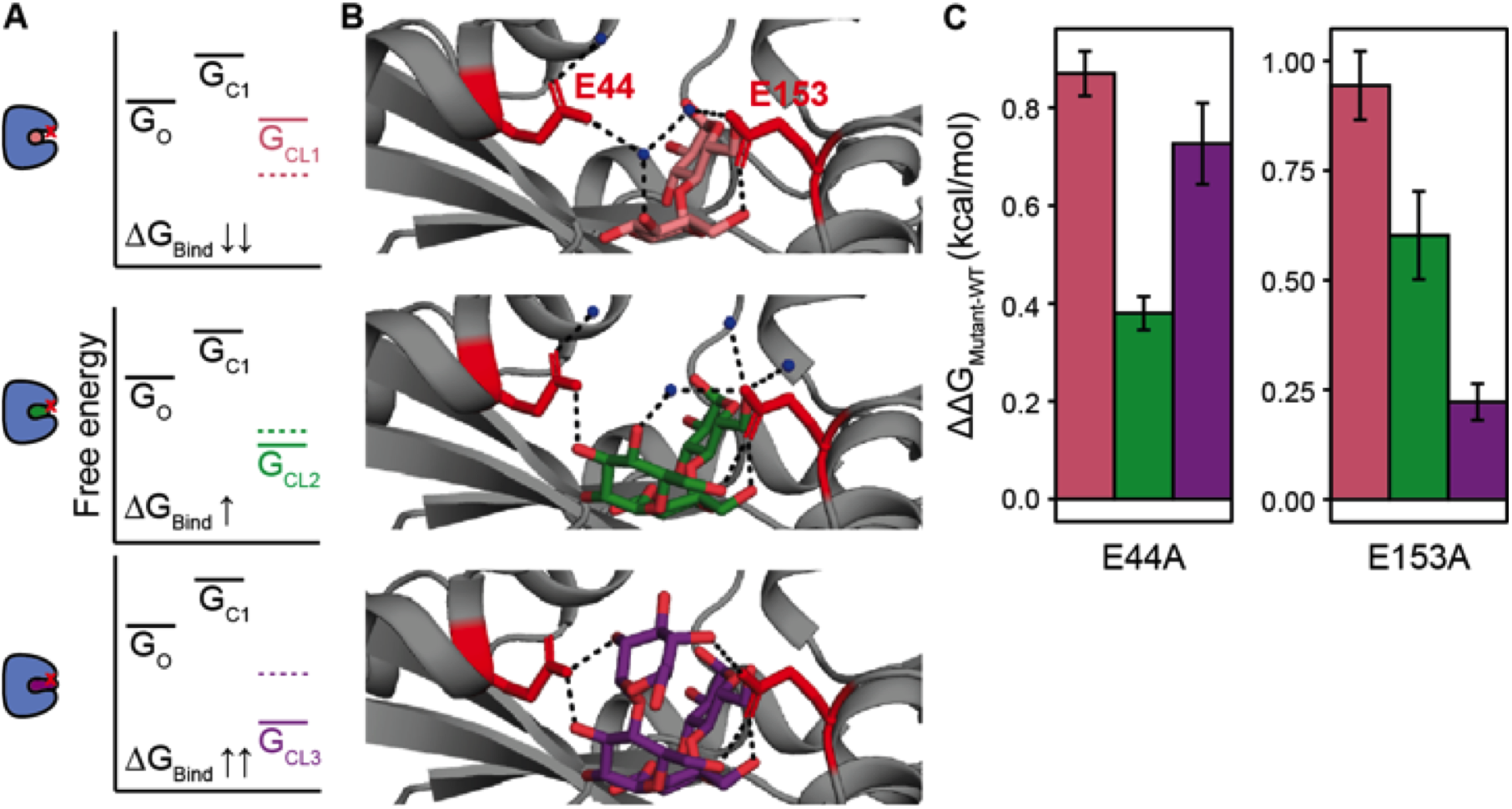
Peripheral binding pocket mutations in MBP exhibit a closed conformation independent ΔΔG_WT-mutant_ between the open and ligand-bound MBP conformation for maltose (pink), maltotriose (green) and maltotetraose (purple). (A) Model to illustrate the idea that binding pocket mutations (red cross in schematic) influence the Δ*G*_bind_, and thereby alter the ligand-bound closed states (G_CL1-3_) independent of the associated closed conformation. The model considers one open state in the absence of ligands (G_O_), one theoretical ligand-free closed state (G_C1_) and three ligand-bound closed states (G_CL1-3_). (B) Zoom-in on crystal structures of MBP bound to maltose (PDB: 1ANF), maltotriose (PDB: 3MBP) or maltotetraose (PDB: 4MBP), respectively. Mutated residues are indicated in red. (C) ΔΔG_WT-mutant_ between the open and ligand-bound MBP conformation for maltose, maltotriose and maltotetraose. Error bars indicate two times the standard error of the mean.

In both E44A and E153A, the maltose, maltotriose and maltotetraose bound conformations are similar to each other, *i.e*. based on the smFRET measurements (**Fig. S2**). As predicted, both mutants exhibit a conformation independent ΔΔG between the open and ligand-bound MBP conformation for the three ligands, which corresponds to changes in K_D_ of a factor between two and five (**Fig. 5C**)(**Table S2**).

## DISCUSSION

Interest in the molecular basis for ligand affinity have grown in recent decades, especially now it is possible to test models more deeply by examining proteins at the level of individual molecules (2, 4, 21, 26). We are no longer limited to static high-resolution structures but can also probe the conformational dynamics of a protein, for instance by single-molecule FRET. The simplest but somewhat naive way to explain ligand affinity deals with the architecture of the binding pocket, which needs to allow for enough protein-ligand interactions and proper orientation of the ligand (4, 13, 43, 44). Differences in ligand affinity of homologous proteins have been explained by differences in binding pocket organization (38, 45). We now provide a thermodynamic framework for analyzing changes in ligand affinity of proteins with mutations in the periphery of or far from the binding pocket. In the case of mutations that are distant from the binding pocket, we rationalize that they can differently affect the free energies of the different ligand-induced closed conformations (**Fig. 2**). We confirm this by showing that these amino acid substitutions induce affinity changes that correlate with the ligand-induced conformation (**Fig. 3**). For mutations in the periphery of the binding pocket we find that the affinity changes do not correlate, because these mutations primarily alter the intrinsic affinity of the binding site for the ligand (Δ*G*_bind_) (**Fig. 5**).

SBPs studied to date bind their ligands via an ‘induced fit’ mechanism (20, 29). For wildtype MBP this means that all ligands bind to a single apo-conformation of the protein, but binding can trigger different closed conformations. Affinity in this case depends on the free energy difference between the open and closed-liganded conformation. Changing the free energy difference between the open and the maltose-induced closed conformation by a single mutation has shown to be sufficient to change the affinity for the ligand by an order of magnitude (22). In this study, we further developed the theory of closed-conformation dependent ligand affinity tuning by describing how single mutations outside the binding pocket can affect the stability of ligand-induced closed MBP conformations. For this, we used the knowledge that maltose, maltotriose and maltotetraose binding induces the same closed conformation in MBP, whereas maltotriitol and β-cyclodextrin trigger two different closed conformations (**Fig. 2C**). Consequently, if a mutation differentially changes the free energy of the three closed conformations, the affinity for maltose, maltotriose and maltotetraose should change by an equal factor, whereas the affinity for maltotriitol and β-cyclodextrin should change differently. The mutations P298G, I317V, N332A and P334G all show the postulated change in affinity (K_D_) for the ligands, *i.e.*, in a manner that correlates with the conformation they induce (**Fig. 3C)**. In the case of P298G, it was even possible to increase the affinity for β-cyclodextrin while decreasing the affinity for maltose, maltotriose, maltotetrasose and maltotriitol.

In nature, evolution could act on the principle of closed conformation-dependent ligand affinity tuning. From an evolutionary point of view, it can be beneficial to use one promiscuous substrate-binding protein like MBP over several specialized homologues (or analogues) when living in nutrient-rich environments (46). However, environments also change in ligand availability on longer timescales and species have to adapt to it. It can then be necessary to evolve the affinity for different ligands according to the need of the species and the availability of the ligand. Selectively tuning of affinity would be hard if it was only restricted to direct protein-ligand interactions. Therefore, tuning of ligand affinities by selectively changing the stability of closed conformations will contribute to this evolutionary need.

Regarding ABC-importer associated SBPs like MBP, several examples of proteins exists for which no ligand could be identified despite clear homology to proteins with known ligand specificity. Two examples are the SBPs YehZ from *E. coli* and BilE from *L. monocytogenes* (47, 48). The proteins are homologous to SBPs that are involved in the binding and transport of quaternary ammonium compounds (QACs). Based on crystal structures, both proteins possess a binding pocket that is very similar to that of other structurally characterized members of this family (47, 48). Yet, the affinity of YehZ and BilE for QACs was at best a 1000-fold lower and no other ligands could be identified (48). It has been shown for homologous OpuAC proteins that different QACs induce different closed conformations (29, 49). Like we show here for MBP, it could well be that amino acid variations outside the binding pocket in YehZ and BilE induce an increase in the Δ*G_conf_* for known QAC-associated closed conformations, while favoring a conformation that is induced by a yet unknown ligand (**Fig. 1**). In favor of this hypothesis, it has been shown that a single non-binding pocket amino acid substitution in YehZ induces an increase in the affinity by an order of magnitude for the QAC glycine betaine (48).

Based on our smFRET measurements on P334G, the maltotetraose-induced conformation had an E* difference of 0.03 when compared to the maltose and maltotriose induced conformations (**Fig. S1D**). Nonetheless, the mutation resulted in a ΔΔG of 0.8 kcal/mol between the open and the closed-liganded conformations of all three ligands (**Fig 3C**). This indicates that the conformational change probably didn’t significantly alter the free energy of the maltotetraose-induced closed conformation compared to the closed conformation that is induced by either maltose or maltotriose. This can be rationalized because P334G and the fluorophore labeling position S352C lie at the opposite ends of a helix that makes direct contacts with maltotetraose through Y341, but not with maltose or maltotriose. Therefore, the structural change of ~0.1 nm is probably very local and not affecting the rest of the protein. Consequently, the effect on the free energy of the conformation is minimal. On the contrary, the mutation S233W was like the other mutations remote from both fluorophore labeling positions but still induced an E* difference of 0.03 and 0.05 between the maltose and maltotriose bound conformation, and maltose and maltotetraose bound conformation, respectively. Because our smFRET measurements provide only spatial information in one dimension, E* differences in this case may reflect more global structural rearrangements that will influence the free energy of the newly formed closed conformations. In this case we can also not exclude possible effects on the Δ*G*_bind_, because the binding pocket architecture could be affected as well. Nonetheless, the mutation S233W shows that it is also possible with only a single mutation to get different closed conformations for maltose, maltotriose and maltotetraose. We also show that the ligands don’t share the same ΔG_conf_ if they induce a different closed conformation. Hence, we speculate that S233W in combination with one of the other mutations allows for uncorrelated affinity tuning of maltose, maltotriose and maltotetraose as well.

We expect that the model we verified here for MBP is not limited to ligand binding proteins but can be applied to enzymes, offering possibilities for protein engineering. Currently, altering the conformational dynamics of an enzyme is a popular tool to engineer its activity. For instance, by changing the availability of catalytically competent conformations (2–4, 6, 9). However, also enzymes adopt different conformations to catalyze different substrates. Examples of such enzymes are the aromatic prenyltransferase AtaPT from *Aspergillus terreus* (43) and rabbit aldolase A (50). It would be valuable to test if our theory can be applied in the optimization of such enzymes. Thereby, we could expand the approaches to evolve enzymes towards biotechnological relevant functions. Moreover, our theory could be of value in understanding and guiding the binding behavior of *de novo* designed proteins (51). Especially, now the existence of multiple conformations are also taken into account during the design process (52).

Although we were able to tune ligand affinity in a conformation-dependent fashion in MBP, it was not possible to reliably predict the magnitude and direction of change in affinity. Therefore, application of our theory still requires some trial and error in finding the most optimal mutations. Furthermore, we intentionally didn’t design mutations that would change the ligand affinities too much, because we were afraid to introduce intrinsic closing in the protein. When MBP would close intrinsically one cannot exclude a possible contribution of ‘conformational selection’ of the ligands. The effect of intrinsic closing/opening dynamics on ligand affinity has been tested for MBP and we intentionally wanted to prevent any contribution of this effect to the affinity changes we were measuring (20, 21). For enzyme engineering studies this is less of a problem if the mutation still leads to an enhancement of the desired activity.

In summary, we present a thermodynamic framework to rationalize changes in ligand affinity by amino acid substitutions distant from the ligand-binding site of the protein, which has been examined for the maltose-binding protein from *E. coli*. We show that ligand affinity can be tuned dependent on the closed conformation that the ligand induces. The new insights help us to understand the sometimes-surprising differences in ligand affinity between homologous proteins and may be used for a more rational optimization of biocatalysts.

## MATERIALS AND METHODS

### Theory

The process of ligand binding to proteins can be treated within the context of Gibbs ensembles (53). The grand partition function Ω of a protein like MBP is

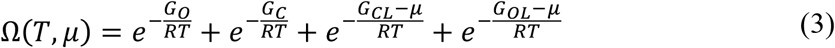

where *R* is the gas constant, *T* is the absolute temperature, *G_i_* is the free energy of state *i* (*O* is the open-unliganded state, *C* is the closed-unliganded state, *OL* is the open-liganded state and *CL* is the closed-liganded state) and *μ* is the chemical potential. We assume that the ligand solution can be treated as an ideal solution, so that *μ* = *μ_0_* + *RT* In (*L*), where *L* is the ligand concentration (relative to 1 Molar) and *μ_0_* is the standard chemical potential (*μ* = *μ_0_* when *L* = 1).

For most SBPs we have that 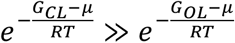 and 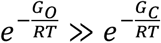 so that **Eq. 3** becomes (29):

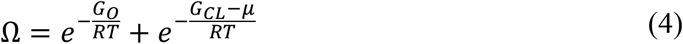

In the presence of ligand, the fraction of proteins occupied by a ligand is

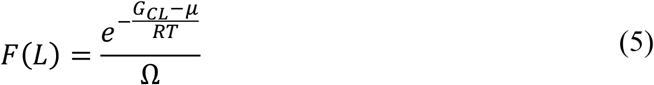

By treating *μ* as an ideal ligand solution 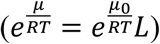 we find that **Eq. 5** is equal to

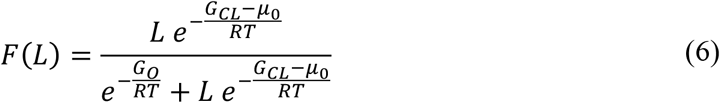

**Eq. 6** can be expressed as the Hill-Langmuir equation, i.e.,

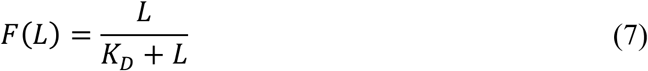

where *K_D_* is the dissociation constant, which is equal to:

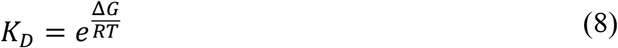

with Δ*G* = *G_CL_* — *µ_0_* — *G_O_*. We can express Δ*G* as

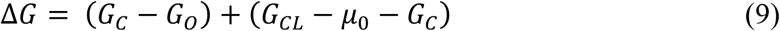

where the conformational free energy is Δ*G_conf_* = *G_C_* — *G_O_* and the intrinsic affinity of the site for the ligand is Δ*G*_bind_ = *G_CL_* — *μ_0_* — *G_C_* so that

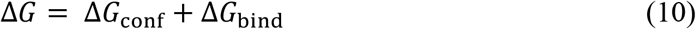

Consequently, the *K_D_* can be altered by modifying the intra-protein interactions (Δ*G_conf_*) and/or the protein-ligand interactions (Δ*G*_bind_).

To relate a change in the *K_D_* back to a change in the Δ*G*, **Eq. 8** has to be rewritten as

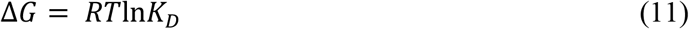

Now, the difference in Δ*G* between a mutant (Δ*G_mutant_*) and wild-type protein (Δ*G_WT_*) binding the same ligand is

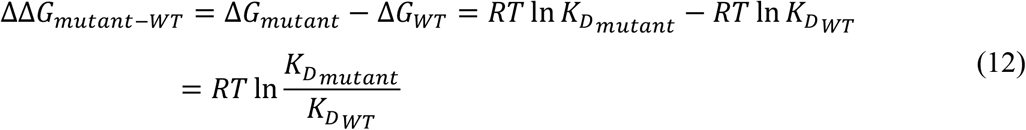

where *K_Dmutant_* and *K_DWT_* are the dissociation constants of the mutant and wild-type protein, respectively. Moreover, we can express ΔΔ*G_mutant-WT_* as

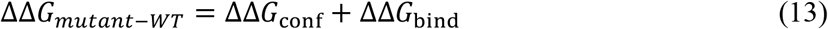

where ΔΔ*G*_conf_ = (Δ*G*_conf_)_*mutant*_ — (Δ*G*_conf_)_*WT*_ and ΔΔ*G*_bind_ = (Δ*G*_bind_)_*mutant*_ — (Δ*G*_bind_)_*WT*_, with (Δ*G*_conf_)_*mutant*_ and (Δ*G*_conf_)_*WT*_ being Δ*G_conf_* of the mutant and wild-type protein, respectively, and (Δ*G*_bind_)_*mutant*_ and (Δ*G*_bind_)_*WT*_ being Δ*G*_bind_ of the mutant and wild-type protein, respectively.

### Materials and culture conditions

All oligonucleotide primers used are listed in the Supplemental Information (**Table S1**). The parental plasmid to introduce mutations in the *malE* gene was pSMB101. pSMB101 originates from the pET20b; the introduced *malE* gene lacks the nucleotides that specify the signal sequence, carries a sequence for a N-terminal His6-tag for protein purification and has the T36C/S352C double mutation for fluorophore labeling of MBP (29). *E. coli* strain MC1061 was used for the amplification and isolation of plasmids. Strain BL21(DE3) was used for expression of the MBP variants. Cells were grown at 37 °C in shaking cultures with Luria–Bertani (LB) medium (1% bactotrypton, 0.5% yeast extract, 1% NaCl) or on plates of LB, supplemented with 1.5% agar and ampicillin (100 μg/mL) for selection. Optical density was monitored at 600 nm (OD600).

The general protein purification buffer consisted of 50 mM Tris-HCl (pH 8.0), 10% glycerol plus 1 mM β-mercaptoethanol, and was supplemented with 1M KCl plus 10 mM imidazole (Buffer A), 50 mM KCl plus 20 mM imidazole (Buffer B) or 50 mM KCl plus 500 mM imidazole (Buffer C). The protein dialysis buffer consisted of 50 mM Tris-HCl (pH 8.0), 50 mM KCl plus 1 mM dithiothreitol (DTT) (Buffer D), which was supplemented with 50% (v/v) glycerol for a second dialysis step (Buffer E).

Maltose and maltotriose were purchased from Sigma-Aldrich. Maltotetraose, malotriitol and β-cyclodextrin were purchased from Biosynth Carbosynth. All maltodextrins were dissolved in 50 mM Tris-HCl (pH 7.4) plus 50 mM KCl (Buffer F).

### Construction of plasmids

pSMB101 was used both as template for the generation of mutants and as plasmid to produce wild type MBP. Site-directed mutagenesis was performed using QuikChange mutagenesis (54). The sequence of each MBP variant was verified by Sanger sequencing (Eurofins Genomics; Mix2Seq).

### Protein expression and purification

Cells harboring a plasmid with one of the MBP variants were grown to 0.5-0.7 OD_600_ before MBP expression was induced by the addition of 250 μM isopropyl β-D-1-thiogalactopyranoside (IPTG). Two hours post-induction, the cells were harvested by centrifugation at 6.000 xg for 10 min at 4 °C. The pellet was washed with ice-cold 50 mM Tris-HCl (pH 8.0) and resuspended in buffer A. Cells were lysed by sonication in the presence of DNase, fresh 1 mM β-mercaptoethanol and 1 mM PMSF. After centrifugation at 27.000 xg for 30 min at 4 °C the supernatant was incubated with 4 mL Ni^2+^-Sepharose resin (GE Healthcare) for 1 hour at 4 °C under gentle agitation. Subsequently, the Ni^2+^-Sepharose was washed with 5 column volumes (CV) of buffer A and 15 CV of buffer B. The His-tagged MBP variants were eluted in buffer C. Absolute protein concentrations were determined, using UV-spectra and an extinction coefficient at 280 nm of 66350 M^−1^cm^−1^ (wildtype and most mutants) or 71850 M^−1^cm^−1^ (MBP variant S233W). The eluted proteins were diluted to 5 mg/mL protein in buffer C supplemented with 5 mM EDTA, dialyzed for 3 hours at 4 °C in 100-400 volumes of buffer D, followed by an overnight dialysis at 4°C in 100-400 volumes of buffer E. The dialyzed solutions were aliquoted and stored at −20 °C. Protein purity was checked on a 12% SDS-PAA gels.

### Protein labeling

A previously described labeling protocol was used to stochastically label the proteins with the two maleimide fluorophores Alexa555 and Alexa647 (Thermofisher Scientific)(19), with some modifications as is described below.

The protein was diluted to 0.5-2 mg/mL in 100 μL buffer F at 4 °C, supplemented with 10 mM DTT in order to reduce all available cysteines. After 10 minutes incubation with 10 mM DTT, the protein was diluted 10 times in buffer F, bound to 90 μL Ni^2+^-sepharose resin and washed twice with 20 CV of buffer F. The fluorophores (50 nanomoles in powder form) were dissolved together in 5 μL of DMSO and 200 times diluted in buffer F. Immediately, they were added to the resin and incubated overnight at 4 °C under gentle agitation; the fluorophore-to-protein ratio was 1:10-50 for both Alexa555 and Alexa647. The resin was washed with 20 CV of buffer F and the protein was eluted in one fraction using buffer C without β-mercaptoethanol and with only 5% glycerol.

### Solution-based smFRET and ALEX

Free fluorophore rotation at positions 36 and 352 in MBP was verified previously by means of steady-state anisotropy (29). Solution-based smFRET and alternating laser excitation (ALEX)(39) experiments were carried out at 5-25 pM of labelled protein at room temperature in buffer A supplemented with additional reagents as stated in the text. Microscope cover slides (no. 1.5H precision cover slides, VWR Marienfeld) were coated with 1 mg/mL BSA for 30-120 seconds to prevent fluorophore and/or protein interactions with the glass material. Excess BSA was subsequently removed by washing and exchange with buffer A. All smFRET experiments were performed using a home-built confocal microscope. In brief, two laser-diodes (Coherent Obis) with emission wavelength of 532 and 637 nm were directly modulated for alternating periods of 50 μs and used for confocal excitation. The laser beams were coupled into a single-mode fibre (PM-S405-XP, Thorlabs) and collimated (MB06, Q-Optics/Linos) before entering an oil immersion objective (60X, NA 1.35, UPlanSAPO 60XO, Olympus). The fluorescence was collected by excitation at a depth of 20 μm. Average laser powers were 30 μW at 532 nm (~30 kW/cm^2^) and 15 μW at 637 nm (~15 kW/cm^2^). Excitation and emission light was separated by a dichroic beam splitter (zt532/642rpc, AHF Analysentechnik), which is mounted in an inverse microscope body (IX71, Olympus). Emitted light was focused onto a 50 μm pinhole and spectrally separated (640DCXR, AHF Analysentechnik) onto two single-photon avalanche diodes (TAU-SPADs-100, Picoquant) with appropriate spectral filtering (donor channel: HC582/75; acceptor channel: Edge Basic 647LP; AHF Analysentechnik). Registration of photon arrival times and alternation of the lasers was controlled by an NI-Card (PXI-6602, National Instruments).

Fluorescent bursts were detected with all photon-burst-search (APBS)(55), using parameters M=15, T=500 and L=25 and a threshold of 150 photons per burst. Three relevant photon counts were extracted from the photon arrival times: acceptor-based acceptor emission (F_AA_), donor-based donor emission (F_DD_) and donor-based acceptor emission (F_DA_). These counts were used to calculate the apparent FRET efficiency E* and stoichiometry S of each photon burst. Apparent FRET efficiency was calculated via:

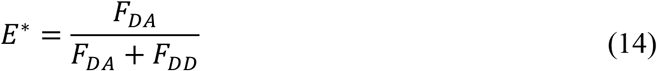

Stoichiometry of a burst was defined as the ratio between overall donor-based emission over the total emission of the burst, i.e.:

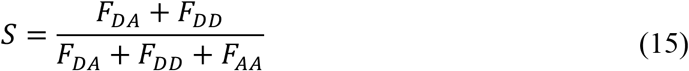

Bursts with a stoichiometry of approximately 0.3-0.7 were selected for further analysis of their apparent FRET efficiency distribution. The apparent FRET efficiency histogram was fitted with a Gaussian distribution, to obtain a 95% confidence interval for the distribution mean (29). Note that, the absolute values of E* can differ significantly between measurements recorded on different days due to drift and optical alignment. Therefore, only measurements recorded on the same day were compared with each other or a standard condition was taken on subsequent days to correct for day-dependent shifts in E*.

### Intrinsic protein fluorescence measurements

Intrinsic protein fluorescence measurements were performed using a Fluorolog-3 spectrofluorometer (Horiba Jobin Yvon). The (unlabeled) proteins were diluted to 0.2 μM in buffer F. The samples were excited at 280 nm (bandwidth: 3 nm) and emission was detected at 365 nm (bandwidth: 3 nm) for β-cyclodextrin/maltotriitol titrations, and at 330 nm (bandwidth: 3 nm) for all other ligands. The signal was measured over 420 seconds and the ligand was titrated at intervals of 20 seconds, starting with the first titration after 20 seconds. To correct for protein dilution and time-dependent bleaching, baselines were recorded by titrating with ligand-free buffer F at the same time intervals. Fluorescence was recorded every 100 ms. The fluorescent signal between ~13-18 seconds after each titration was averaged and used for subsequent analysis. The dissociation constants (K_D_) were calculated by fitting (25, 56):

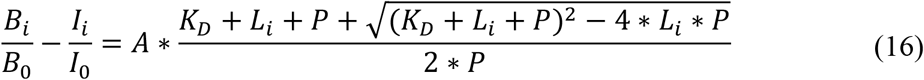

Where I_i_ is the mean fluorescence intensity, B_i_ the mean intensity of the corresponding baseline and L_i_ the ligand concentration at interval i. I_0_ and B_0_ are the mean fluorescence intensities at the first interval. The protein concentration P was fixed at 0.2 μM. Both the asymptote (A) and the K_D_ were defined as free fit parameters and fitted with a nonlinear least squares model in R (nls function from the R Stats Package v.3.6.2). The ΔΔG_WT-mutant_ was calculated via **Eq. 12**. The standard error of the mean (SEM) was calculated via:

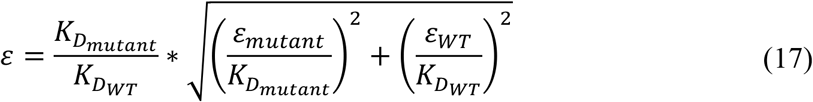

where *K_DWT_*, *ε_WT_*, and *K_Dmutant_* and *ε_mutant_* refer to the dissociation constant and SEM of wild-type, and mutant MBP, respectively.

## AUTHOR CONTRIBUTIONS

B.P. supervised the project. M.v.d.N. performed the molecular biology and fluorescence measurements. M.d.B. and M.v.d.N performed the smFRET measurements. M.d.B developed the theoretical framework. M.v.d.N wrote the initial manuscript. M.d.B., M.v.d.N and B.P. finalized the manuscript. All authors contributed to the discussion and interpretation of the results.

## ACKNOWLEGDEMENTS

This work was financed by the NWO Gravitation Program BaSyC and the European Research Council (ERC Advanced Grant ABCvolume #670578).

## DATA AVAILABILITY

All study data are included in this article and/or Supplementary Information appendix.

## Notes

### Competing Interest Statement

The authors have declared no competing interest.

