## Supplementary data for "Stability of ligand-induced protein conformation influences affinity in maltose-binding protein"

Table S1. Primers used in this study to generate mutations in MBP (T36C/S352C)

| **Oligo nr.** | **Oligo name** | **Sequence (from 5' to 3')** † |
|---|---|---|
| 8152 | E44A-fw | gcatccggataaactgGCAgagaaattcccacaggttgc |
| 8153 | E44A-rev | gcaacctgtgggaatttctcTGCcagtttatccggatgc |
| 8392 | E153A-fw | gatgttcaacctgcaaGCAccgtacttcacctggccg |
| 8393 | E153A-rev | cggccaggtgaagtacggTGCttgcaggttgaacatc |
| 8162 | S233G-fw | ccgtgggcatggGGCaacatcgataccagcaaag |
| 8163 | S233G-rev | ctttgctggtatcgatgttGCCccatgcccacgg |
| 8164 | S233W-fw | ccgtgggcatggTGGaacatcgataccagcaaag |
| 8165 | S233W-rev | ctttgctggtatcgatgttCCAccatgcccacgg |
| 8158 | P298G-fw | ggttaataaagacaaaGGTctgggtgccgtagcg |
| 8159 | P298G-rev | cgctacggcacccagACCtttgtctttattaacc |
| 8156 | I317V-fw | cgaaagatccacgtGTGgccgccactatggaaaacg |
| 8157 | I317V-rev | cgttttccatagtggcggcCACacgtggatctttcg |
| 8402 | N332A-fw | ggtgaaatcatgccgGCTatcccgcagatgtcc |
| 8403 | N332A-rev | ggacatctgcgggatAGCcggcatgatttcacc |
| 8398 | P334G-fw | catgccgaacatcGGTcagatgtccgctttctgg |
| 8399 | P334G-rev | ccagaaagcggacatctgACCgatgttcggcatg |

† Mutated triplets are in capitals

Table S2. Dissociation constants ($K_D$ values) obtained from fluorescence titrations of MBP variants

| **MBP variant** | **Ligand** | **$K_D$ (nM)** | **2x(error of the mean)** | **Replicates** |
|---|---|---|---|---|
| WT | Maltose | 2,140 | 103 | 9 |
| | Maltotriose | 835 | 68 | 11 |
| | Maltotetraose | 554 | 58 | 8 |
| | Maltotriitol | 51,010 | 3,012 | 9 |
| | β-cyclodextrin | 2,019 | 77 | 6 |
| E44A | Maltose | 9,537 | 200 | 3 |
| | Maltotriose | 1,603 | 61 | 4 |
| | Maltotetraose | 1,928 | 89 | 4 |
| E153A | Maltose | 10,828 | 729 | 4 |
| | Maltotriose | 2,348 | 341 | 6 |
| | Maltotetraose | 811 | 125 | 4 |
| S233W | Maltose | 6,162 | 122 | 3 |
| | Maltotriose | 976 | 52 | 4 |
| | Maltotetraose | 420 | 16 | 4 |
| P298G | Maltose | 4,167 | 91 | 3 |
| | Maltotriose | 1,722 | 182 | 3 |
| | Maltotetraose | 1,027 | 57 | 4 |
| | Maltotriitol | 81,345 | 4,085 | 3 |
| | β-cyclodextrin | 1,203 | 39 | 3 |
| I317V | Maltose | 594 | 93 | 5 |
| | Maltotriose | 211 | 16 | 5 |
| | Maltotetraose | 128 | 8 | 4 |
| | Maltotriitol | 5,453 | 165 | 3 |
| | β-cyclodextrin | 319 | 25 | 3 |
| N332A | Maltose | 3,240 | 258 | 4 |
| | Maltotriose | 1,347 | 202 | 4 |
| | Maltotetraose | 954 | 132 | 5 |
| | Maltotriitol | 68,809 | 3,971 | 12 |
| | β-cyclodextrin | 3,843 | 131 | 10 |
| P334G | Maltose | 7,729 | 223 | 3 |
| | Maltotriose | 3,312 | 342 | 4 |
| | Maltotetraose | 2,227 | 100 | 4 |
| | Maltotriitol | 98,798 | 7,718 | 3 |
| | β-cyclodextrin | 3,072 | 87 | 7 |

Table S3. Number of analyzed bursts per MBP variant and condition

| MBP variant | Condition | Number of analyzed bursts |
|---|---|---|
| WT | Ligand free | 3,822 |
| | 1 mM maltose | 4,518 |
| | 1 mM maltotriose | 3,074 |
| | 1 mM maltotetraose | 2,844 |
| | 1 mM maltotriitol | 3,783 |
| | 100 µM β-cyclodextrin | 3,134 |
| E44A | Ligand free | 2,806 |
| | 1 mM maltose | 3,007 |
| | 1 mM maltotriose | 2,900 |
| | 1 mM maltotetraose | 1,618 |
| E153A | Ligand free | 2,455 |
| | 1 mM maltose | 3,518 |
| | 1 mM maltotriose | 4,313 |
| | 1 mM maltotetraose | 3,597 |
| P298G | Ligand free | 2,443 |
| | 1 mM maltose | 2,234 |
| | 1 mM maltotriose | 2,374 |
| | 1 mM maltotetraose | 2,113 |
| | 1 mM maltotriitol | 2,669 |
| | 100 µM β-cyclodextrin | 3,007 |
| S233W | Ligand free | 6,377 |
| | 1 mM maltose | 3,610 |
| | 1 mM maltotriose | 2,141 |
| | 1 mM maltotetraose | 2,096 |
| I317V | Ligand free | 2,368 |
| | 1 mM maltose | 2,080 |
| | 1 mM maltotriose | 1,707 |
| | 1 mM maltotetraose | 2,228 |
| | 1 mM maltotriitol | 2,719 |
| | 100 µM β-cyclodextrin | 4,252 |
| N332A | Ligand free | 2,600 |
| | 1 mM maltose | 803 |
| | 1 mM maltotriose | 1,716 |
| | 1 mM maltotetraose | 1,280 |
| | 1 mM maltotriitol | 2,733 |
| | 100 µM β-cyclodextrin | 2,216 |
| P334G | Ligand free | 2,983 |
| | 1 mM maltose | 2,123 |
| | 1 mM maltotriose | 4,185 |
| | 1 mM maltotetraose | 2,349 |
| | 1 mM maltotriitol | 2,506 |
| | 100 µM β-cyclodextrin | 2,863 |

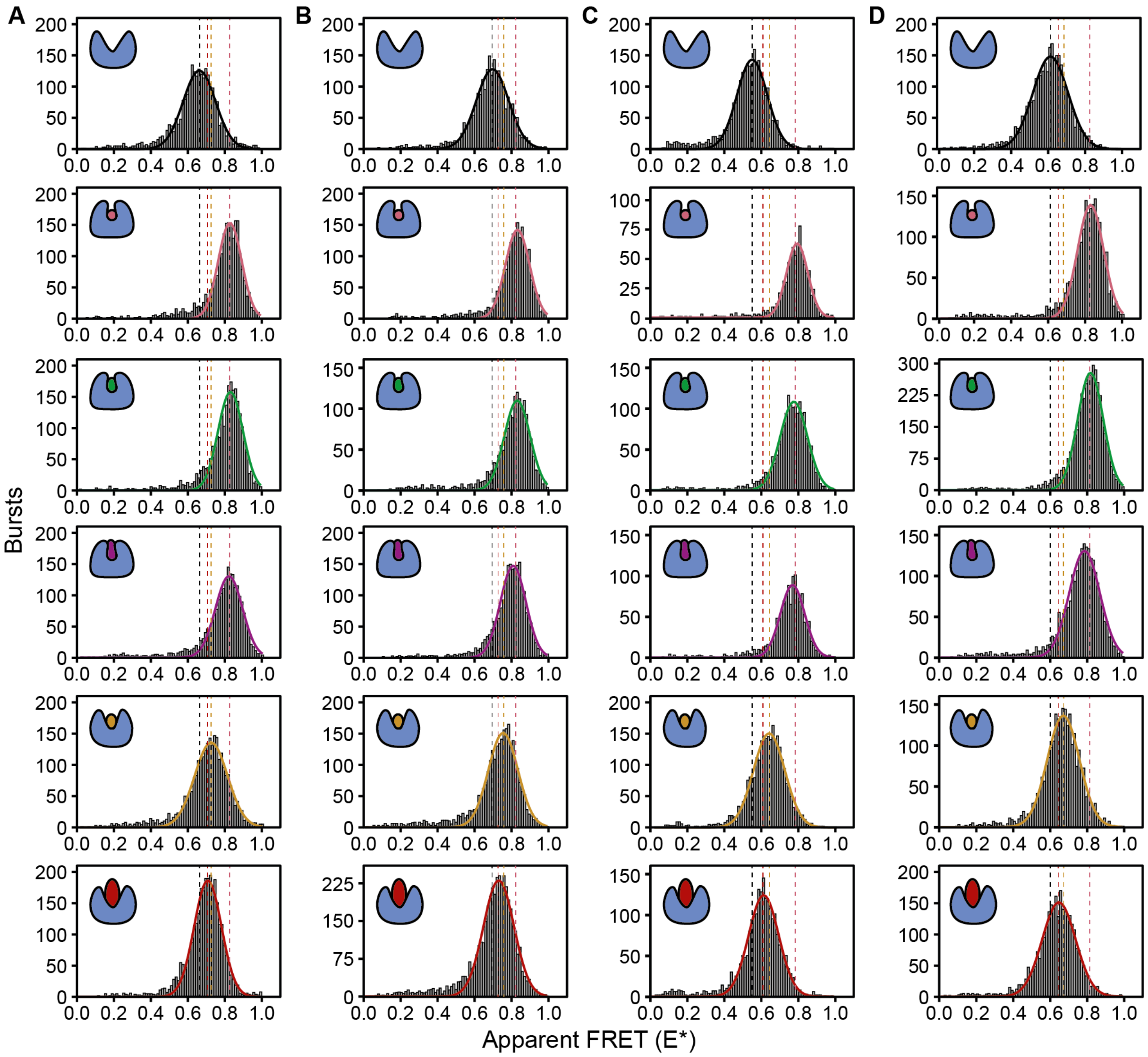


Figure S1. Solution-based smFRET histograms of MBP variants with substitutions distant from the binding pocket: a) P298G, b) I317V, c) N332A and d) P334G in the absence of any ligand; or in the presence of 1 mM maltose (pink), maltotriose (green), maltotetraose (purple), maltotriitol (yellow), or β-cyclodextrin (red). Solid lines indicate best fit of a single Gaussian distribution, as described in Materials and Methods. The associated means are represented as dotted lines. The dotted lines in pink represent a general mean for the conditions with maltose, maltotriose and maltotetraose. Numbers of included bursts per plot are given in **Table S3**.

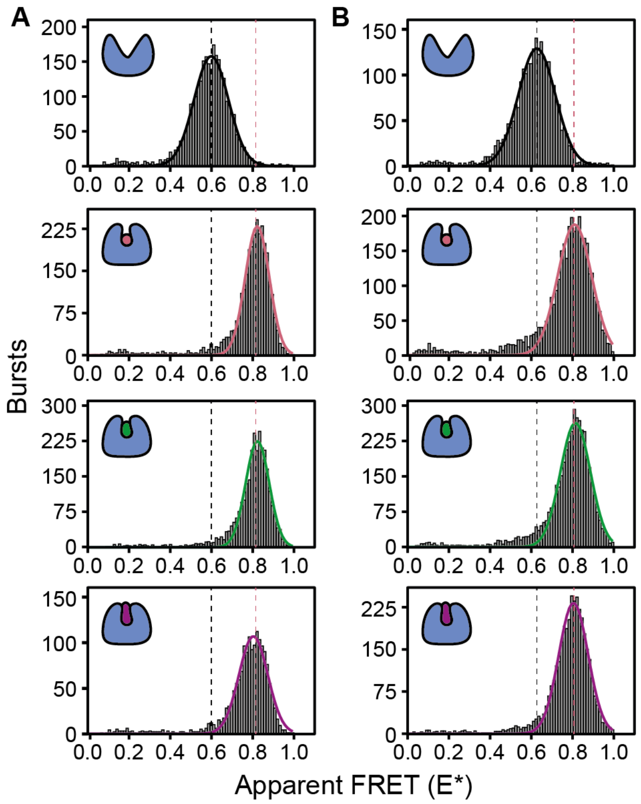

Figure S2. Solution-based smFRET histograms of MBP variants with substitutions in the periphery of the binding-pocket: a) E44A and b) E153A in the absence of any ligand; or in the presence of 1 mM maltose (pink), maltotriose (green), or maltotetraose (purple). Solid lines indicate best fit of a single Gaussian distribution, as described in Materials and Methods. The associated means are represented as dotted lines. The dotted lines in pink represent a general mean for the conditions with maltose, maltotriose and maltotetraose. Numbers of included bursts per plot are given in **Table S3**.
